# Huntingtin inclusion bodies have distinct immunophenotypes and ubiquitination profiles in the Huntington’s disease human cerebral cortex

**DOI:** 10.1101/2025.02.03.636327

**Authors:** Molly E. V. Swanson, Adelie Y. S. Tan, Lynette J. Tippett, Clinton P. Turner, Maurice A. Curtis, Emma L. Scotter, Hilal A. Lashuel, Mike Dragunow, Richard L. M. Faull, Helen C. Murray, Malvindar K. Singh-Bains

**Affiliations:** Centre for Brain Research, University of Auckland, Auckland, New Zealand; School of Biological Sciences, University of Auckland, Auckland, New Zealand; Department of Anatomy and Medical Imaging, University of Auckland, Auckland, New Zealand; Department of Pharmacology and Clinical Pharmacology, University of Auckland, Auckland, New Zealand; School of Psychology, University of Auckland, Auckland, New Zealand; Department of Anatomical Pathology, LabPlus, Auckland City Hospital, Auckland, New Zealand; Laboratory of Molecular and Chemical Biology of Neurodegeneration, Brain Mind Institute, Ecole Polytechnique Federale de Lausanne (EPFL), Lousanne, Switzerland

## Abstract

Huntington’s disease (HD) is a hereditary neurodegenerative condition caused by a CAG repeat expansion mutation in the gene encoding the huntingtin (Htt) protein. The accumulation of Htt inclusion bodies is a pathological hallmark of HD and a common target for therapeutic strategies. However, the limited efficacy of treatments targeting the Htt protein highlights the need for a better understanding of the role of Htt inclusion bodies in HD pathogenesis. This study examined the heterogeneity of Htt inclusion body composition by co-labelling with three Htt epitope-specific antibodies to characterize Htt inclusion body ‘immunophenotype’. We then characterized the size and sub-cellular location of Htt inclusions with distinct immunophenotypes. Using multiplex immunohistochemistry, we also examined the ubiquitination profile of each immunophenotype. Our findings demonstrate that Htt inclusions have a range of immunophenotypes, with some labelled by only one of the three antibodies and others exhibiting co-labelling by several antibodies, thus demonstrating the heterogeneity in inclusion composition and structure. We outline evidence that inclusion bodies exclusively labelled with the EM48 antibody are small, non-nuclear, and more abundant in HD cases with increased CAG repeat length, higher Vonsattel grade, and earlier age of onset. We also find that Htt inclusion bodies labelled by multiple antibodies are more likely to be ubiquitinated, predominantly by K63-rather than K48-linked ubiquitin, suggesting preferential degradation by autophagy. Lastly, we show that ubiquitinated Htt inclusion bodies are more highly immunoreactive for ubiquilin 2 than p62. Our findings highlight the need for multiple antibodies to capture the full spectrum of Htt pathology in HD and imply that future studies should consider the diversity of inclusion body composition and structure when correlating pathology formation to neurodegeneration, clinical symptoms, or disease severity.

## Introduction

Huntington’s disease (HD) is a hereditary neurodegenerative condition characterized by a triad of motor, mood, and cognitive symptoms (1). HD is caused by an autosomal dominantly inherited cytosine-adenine-guanine (CAG) trinucleotide repeat expansion within exon 1 of the huntingtin gene (HTT) on chromosome 4, encoding the huntingtin (Htt) protein (2). The misfolding of mutant Htt protein within the brain, evidenced by the deposition of Htt aggregates, is hypothesized to trigger pathogenic processes causing progressive neuronal cell death, affecting the basal ganglia, cerebral cortex and cerebellum (3–10). However, there is still controversy over the relative contributions of both normal and mutant Htt to the pathogenesis of HD. The development of effective tools to study Htt protein and capture the diversity of its proteoform and pathological diversity within the human brain is critical to inform the development of disease-modifying treatments (11, 12).

The ability to identify and phenotype normal and mutant Htt in the human brain has primarily been limited by the available antibodies (3, 13); the most common antibodies used to profile Htt inclusion bodies (1C2, 1F8, S830, and EM48) are limited by epitope specificity and/or commercial availability. Furthermore, with the biochemical and structural diversity of Htt inclusion bodies, it is unlikely that any single antibody is capable of immunolabelling all pathogenic Htt, as recently demonstrated for misfolded proteins linked to other neurodegenerative diseases (14–16). Supporting this hypothesis, we recently identified that antibodies targeting different Htt epitopes show different patterns of immunoreactivity in the HD brain (12). Three commercially available antibodies directed to the Htt N-terminus, MAB5374 (EM48), EPR5526, and MW1, successfully immunolabelled Htt inclusion bodies in HD, but each labelled a different number of aggregates (12). It remains unclear whether these antibodies are identifying distinct Htt inclusion bodies or inclusion bodies with overlapping composition and structure A deeper understanding of Htt inclusion body phenotypes will aid in the development in disease-modifying treatments targeting mutant Htt.

Until more recently, studies which have focused on correlating Htt pathology burden and neurodegeneration and disease stages or severity have treated all aggregates as if they are equal, despite the large morphological diversity of Htt pathology in post-mortem human brain tissue. Recent studies on alpha synuclein pathology suggest that post translation modifications (PTMs) play a central role at different stages of alpha synuclein pathology formation and maturation of Lewy bodies. For example, it has been recently shown that the presence of specific PTMs on the surfaces of fibrils, such as O-Glc-NAC modifications or nitration, nearly abolishes the seeding activity of alpha synuclein aggregates (17). Htt undergoes a large variety of PTMs including phosphorylation (18), SUMOylation (19), acetylation (20), proteolysis (21), palmitoylation (22) and ubiquitination (23). Therefore, the differences in the structural and biochemical properties of protein aggregates in neurodegenerative diseases could underly not only their morphological diversity but also their toxicity and pathogenic properties. If true, this could have significant implications for developing antibody-based therapies and diagnostics for HD and other neurodegenerative diseases.

Ubiquitination involves the tagging of a target protein with chains of ubiquitin with specific ‘linkages’ for targeting to specific cellular pathways, including protein degradation (24). Tetraubiquitin chains in which each ubiquitin is linked to the next ubiquitin via internal lysine residue K48 (K48-linked) predominantly target cargo to the proteasome, while K63-linked ubiquitin chains promote aggregation and predominantly target cargo to autophagy (25, 26). Indeed, K48-versus K63-linked ubiquitination of Htt inclusion bodies in a mouse model of HD promoted degradation versus aggregation, respectively (27). An age-dependent shift in Htt ubiquitination from K48-linked to K63-linked is hypothesised to contribute to late-onset neuropathology in HD (27). Therefore, these differential ubiquitination patterns of Htt may suggest a cellular attempt to degrade aggregates and potential defects with the processing or clearance of ubiquitinated Htt aggregates. While Htt inclusion bodies in the human HD brain are known to be ubiquitinated (13, 28), the specific ubiquitin linkages and how this relates to inclusion composition and structure remain unknown.

Ubiquitination targets proteins to UPS or autophagy through interaction with triage proteins, including ubiquilin 2 and p62. Ubiquilin 2 plays a significant role in the UPS; its ubiquitin-associated domain enables it to associate with ubiquitinated cargo and its ubiquitin-like domain mediates interactions with the proteosome (29). However, ubiquilin 2 can also interact with microtubule-associated protein 1 light chain 3 (LC3), a key component of autophagosome membranes, implicating ubiquilin 2 in autophagy (30). P62 was initially thought to be solely involved in autophagy (31, 32), with a ubiquitin-associated domain to interact with K63-linked polyubiquitin chains and an LC3 interacting region (LIR) domain for direct binding to LC3 to direct substrates to autophagosomes (32, 33). However, p62 also binds to proteasome complexes through its Phox and Bem1 (PB1) domain, facilitating the shuttling of ubiquitinated proteins bound at its ubiquitin-like domain to the proteosome (34, 35). Therefore, while ubiquilin 2 and p62 cannot be used to assess specific recruitment to either the UPS or autophagy, their interaction with ubiquitinated proteins suggests preferential degradation by one of these two pathways. Interestingly, the interaction of Htt inclusion bodies with ubiquilin 2 and p62 in the human HD brain remains largely unknown.

Therefore, this study aims to investigate the ubiquitination profile for three key Htt inclusion immunophenotypes previously validated to be HD-specific in human brain tissue microarrays (12). Towards this goal, the ubiquitination profile was examined using multiplex immunohistochemical approaches on cortical human brain tissue microarrays comprising 55 human tissue samples from HD and neurologically normal control cases, with a focus on key ubiquitin linkages which target proteins for ubiquitination, including K48 (UPS-targeting) versus K63 (promotes autophagy), and triage proteins, ubiquilin 2 and p62. Furthermore, we assessed the size and spatial localisation of the profiled inclusions to better understand the heterogeneity of Htt aggregates. We hypothesized that this approach would contribute to identifying pathogenic Htt species based on their ubiquitination profiles, both aiding in understanding the biochemical features underlying the diversity of Htt pathology and helping guide the development of therapeutic approaches to target and clear pathogenic Htt.

## Methods

### Human Brain Tissue

Post-mortem human brain tissue was provided by the Neurological Foundation Human Brain Bank at the University of Auckland, Centre for Brain Research. The tissue was donated with informed consent from the donor and family before brain removal. All procedures and protocols were approved by the Health and Disability Ethics Committee (Ref.14/NTA/208) and all research presented in this study was performed in accordance with the approved guidelines and regulations.

Fixation and dissection protocols for human brain tissue preparation have been described previously (36, 37). Briefly, the brains were perfused with 15% formaldehyde in 0.1 M phosphate buffer through the cerebral arteries. For each case, a 5-mm thick slice of the middle temporal gyrus (MTG) was embedded in paraffin wax to be used to construct tissue microarrays (TMAs). The MTG has previously been implicated in HD pathogenesis, with a demonstrated reduction in SMI-32+ pyramidal neurons in HD compared to the normal control brains, confirming neuronal cell loss and thereby justifying the use of this region for further HD neuropathology studies (6). The TMAs consisted of post-mortem brain tissue from the MTG of 27 neurologically normal control cases and 28 HD cases (Table 1). The normal and HD cases were matched as closely as possible for age at death, sex, and post-mortem delay (PMD).

**Table 1.** Characteristics of Huntington’s disease and normal cases utilized for this study (TMA 18C).

| Case Number | Diagnosis | Sex | CAG repeat length | Age of Death (y) | PMD (h) | Cause of Death |
| --- | --- | --- | --- | --- | --- | --- |
| H118 | Normal | M | 15/16 | 57 | 10 | Coronary artery disease |
| H120 | Normal | M | 18/22 | 62 | 11 | Ischemic heart disease |
| H127 | Normal | F | 15/17 | 59 | 21 | Pulmonary embolism |
| H129 | Normal | M | 20/21 | 48 | 12 | Pulmonary embolism |
| H130 | Normal | M | 17/22 | 32 | 13 | Pulmonary embolism following surgery |
| H131 | Normal | F | 17/17 | 73 | 13 | Ischemic heart disease |
| H136 | Normal | M | N/A | 75 | 13 | Ruptured abdominal aortic aneurysm |
| H139 | Normal | M | N/A | 73 | 5.5 | Ischemic heart disease |
| H183 | Normal | M | 17/17 | 61 | 13 | Multiple-organ failure |
| H193 | Normal | M | 17/24 | 71 | 23 | Valvular heart disease coronary atherosclerosis |
| H194 | Normal | M | 17/19 | 68 | 22.5 | Coronary atherosclerosis |
| H195 | Normal | M | 18/18 | 65 | 18 | Ischemic heart disease |
| H198 | Normal | F | 17/23 | 67 | 27 | Acute pyelonephritis |
| H200 | Normal | M | 16/17 | 56 | 23 | Asphyxia |
| H202 | Normal | M | 19/24 | 83 | 14 | Ruptured abdominal aortic aneurysm |
| H204 | Normal | M | 18/18 | 66 | 9 | Ischemic heart disease |
| H211 | Normal | M | 17/18 | 41 | 8 | Ischemic heart disease |
| H215 | Normal | F | 17/17 | 67 | 23.5 | Ischemic heart disease |
| H230 | Normal | F | N/A | 57 | 32 | Carcinomatosis (renal) |
| H231 | Normal | M | N/A | 65 | 8 | Ischemic heart disease |
| H238 | Normal | F | 14/16 | 63 | 16 | Dissecting aortic aneurysm |
| H239 | Normal | M | 13/15 | 64 | 15.5 | Ischemic heart disease |
| H243 | Normal | F | 17/17 | 77 | 13 | Coronary atherosclerosis |
| H245 | Normal | M | 21/24 | 63 | 20 | Asphyxia |
| H246 | Normal | M | 17/18 | 88 | 17 | Type II myocardial infarction |
| H252 | Normal | F | 18/19 | 94 | 13.5 | Lower respiratory tract infection; congestive heart failure |
| H253 | Normal | M | 16/17 | 96 | 24 | Deconditioning |
| HC051 | HD | M | 16/43 | 58 | 4.5 | Pneumonia |
| HC052 | HD | F | 19/46 | 61 | 23 | Cerebrovascular accident |
| HC075 | HD | M | 18/52 | 41 | 5.5 | Pneumonia |
| HC076 | HD | M | 19/42 | 71 | 16 | Pneumonia |
| HC088 | HD | F | 17/44 | 57 | 19 | Myocardial infarction |
| HC089 | HD | F | 17/41 | 71 | 12 | Pneumonia |
| HC091 | HD | F | 23/44 | 53 | 5.5 | HD |
| HC096 | HD | F | 22/53 | 39 | 20 | Starvation with dehydration |
| HC102 | HD | M | 17/42 | 64 | 10 | Aspiration pneumonia |
| HC104 | HD | M | 18/51 | 40 | 15 | Pneumonia |
| HC119 | HD | M | 17/48 | 51 | 15.5 | Dehydration |
| HC123 | HD | M | 23/47 | 54 | 14 | Bronchopneumonia |
| HC128 | HD | M | N/A | 80 | 13 | Myocardial infarction |
| HC133 | HD | M | 17/43 | 65 | 13 | Renal failure |
| HC134 | HD | M | 18/43 | 62 | 9 | HD |
| HC137 | HD | M | 17/41 | 83 | 13 | Pneumonia |
| HC140 | HD | M | 17/40 | 62 | 22 | Chronic renal failure |
| HC141 | HD | F | -/42 | 84 | 14 | N/A |
| HC142 | HD | F | 15/45 | 55 | 25 | Pneumonia |
| HC143 | HD | F | 17/54 | 45 | 16 | Respiratory arrest |
| HC144 | HD | M | 10/46 | 55 | N/A | Leukaemia |
| HC145 | HD | M | N/A | 54 | N/A | Chest infection |
| HC146 | HD | M | 17/47 | 46 | 6 | HD |
| HC147 | HD | M | 27/42 | 64 | 18 | Pulmonary thromboembolus |
| HC148 | HD | M | 22/43 | 63 | 16 | Pulmonary embolism |
| HC150 | HD | F | 22/42 | 64 | 21 | Bronchopneumonia |
| HC151 | HD | M | 17/43 | 71 | 35.5 | Pre-renal failure, poor fluid intake<br>and swallowing |
| HC152 | HD | M | 23/43 | 60 | 16 | Renal failure |

Abbreviations: F, female; HD, Huntington's disease; M, male; N/A, not available; PMD, post-mortem delay

The normal cases included in the TMAs (Table 1) consisted of 19 males and 8 females, with an age at death of 32–96 years [mean = 66.3 ± 14.0 years (mean ± SD)], and PMD of 8–24 h (mean = 16.2 ± 6.4 h). The HD cases included in the TMAs consisted of 18 males and 10 females, with an age at death of 39–84 years (mean = 59.7 ± 11.7 years), and PMD of 4.5–35.5 h (mean = 15.3 ± 6.7 h). The age at symptom onset for HD cases was 15–70 years (mean = 38.5 ± 12.7 years). The age of clinical onset was based on well-defined criteria outlined by Tippett et al. (2007) (10). For both normal and HD cases, the polyglutamine (CAG) repeat length was determined using polymerase chain reaction from either a blood sample or cerebellar brain tissue sample (Tables 1). The HD cases were neuropathologically graded based on the degree of striatal atrophy according to the Vonsattel grading system (38).

### Tissue microarray (TMA) construction

The human brain TMAs utilized in this study were constructed according to the protocol by Singh-Bains et al. (2021) (39). The TMA utilized in this study consisted of cylindrical cores of formalin-fixed, paraffin-embedded human brain tissue obtained from the MTG of the 27 neurologically normal control and 28 HD cases outlined in Table 1 and Table 2. Prior to TMA construction, a section from each designated donor block of human brain tissue was stained with cresyl violet (Nissl stain) to enable the identification of layers II-V of the cerebral cortex and provide a guide for tissue coring, ensuring only grey matter was captured for each case. An adjacent section was labelled for a ubiquitously expressed antibody, anti-glial fibrillary acidic protein (GFAP), to ensure high-quality antigenicity of the tissue. Subsequently, 2-mm-diameter cores were extracted from each donor block of tissue and inserted in a recipient block of paraffin wax using the Veridium VTA-100 Tissue Microarrayer. Once the coring process was complete, the TMA recipient block was cut into 7-μm-thick sections using a microtome (RM2235, Leica) and mounted onto positively charged slides (Uber Frost) in preparation for fluorescent immunohistochemistry.

**Table 2.** Antibody panels used for immunohistochemistry on HD tissue microarrays.

| Antibody | Manufacturer & Product Number | Host | Dilution | Secondary Antibody | Manufacturer / Product Number | Labelling round |
| --- | --- | --- | --- | --- | --- | --- |
| Ubiquilin 2 | Santa Cruz Sc100612 | Mouse IgG2a | 1:1,000 | Goat anti-mouse IgG2a 488 | ThermoFisher A21131 | 2 |
| K48-linked ubiquitin | Merck 51307 | Rabbit | 1:300 | Goat anti-rabbit 488 | ThermoFisher A11034 | 3 |
| K63-linked ubiquitin | Novus HWA4C4 | Mouse IgG2a | 1:300 | Goat anti-mouse IgG2a 647 | ThermoFisher A21241 | 3 |
| Ubiquitin | ThermoFisher 131600 | Mouse IgG1 | 1:500 | Goat anti-mouse IgG1 594 | ThermoFisher A21123 | 3 |
| P62 | Progen GP62-C | Guinea pig | 1:500 | Goat anti-guinea pig 546 | ThermoFisher A11076 | 3 |
| EM48 | Merck MAB5374 | Mouse IgG1 | 1:200 | Goat anti-mouse IgG1 488 | ThermoFisher A21121 | 5 |
| MW1 | Merck MABN242 | Mouse IgG2b | 1:500 | Goat anti-mouse IgG2b 647 | ThermoFisher A21242 | 5 |
| EPR5526 | Abcam ab109115 | Rabbit | 1:1,000 | Goat rabbit 594 | ThermoFisher A11037 | 5 |

### Multiplex Immunohistochemistry

The images used for the analysis presented in this study were acquired as part of a larger multiplex image dataset. Multiplex fluorescence immunohistochemistry was performed as previously described (40, 41). Six-plex labelling was achieved by combining antibodies from each available IgG subclass of mouse antibody (IgG1, IgG2a, IgG2b, IgG3), with antibodies raised in non-mouse hosts (rat, rabbit, guinea pig, chicken), across five rounds of labelling. The antibodies analysed in this study were used in rounds 3 and 5, as per Table 2. One TMA section was labelled. The section was deparaffinised and subjected to heat-mediated epitope retrieval using 10 mM Tris-EDTA buffer pH 9.0 in a pressure cooker (2100 Antigen Retriever, Aptum Biologics Ltd). TrueBlack reagent (Biotum) was applied as per the manufacturer’s instructions to quench endogenous autofluorescence and 10% normal goat serum diluted in phosphate-buffered saline (PBS) was subsequently applied for 1 hour at room temperature to block non-specific binding of the secondary antibody. The sections were then incubated overnight in the primary antibody cocktail (diluted in 1% normal goat serum in PBS) for round 1 at 4 °C, followed by incubation in the appropriate species-specific secondary antibody cocktail, containing Hoechst to stain nuclei, for 4 h at room temperature. The section was washed in PBS between all steps. The slide was coverslipped with Prolong Gold antifade mountant (ThermoFisher) and imaged on an automated widefield fluorescence microscope (Zeiss Z2 Axioimager) equipped with MetaSystems Metafer and Vslide software, with a 20 × air objective (0.9 NA). This microscope is equipped with 6 custom excitation/dichroic/emission filter sets optimised for spectral separation of compatible fluorophores as previously described (42). The section was then decoverslipped by immersion in PBS and antibodies were stripped from sections by applying 5X NewBlot™ Nitro Stripping Buffer (Li-Cor) undiluted for 10 min at room temperature. The section was washed again in PBS, and a subsequent round of immunostaining and imaging was performed as above. In total, five rounds of labelling were performed. The images from all five rounds were aligned using a custom Python script (41, 43). The images for K48-linked ubiquitin, K63-linked ubiquitin, pan-ubiquitin, EM48, EPR, MW1, p62, and ubiquilin 2 were used for analysis in this study (Table 2).

### Image Analysis

To identify and quantify ubiquitination and triage protein tagging of Htt inclusion bodies, we developed custom image analysis pipelines in Metamorph software (Molecular Devices) similar to those previously described (41, 44–46). Prior to analysis, manual regions of interest (ROI) were drawn on Hoechst images to retain TMA tissue area and exclude tissue folds and defects.

To identify all Htt inclusion bodies, we used an intensity threshold on the EM48, EPR, and MW1 images to create binary masks of EM48+, EPR+, and MW1+ Htt inclusion bodies. Thresholds were set manually for each case to segment positive labelling only. Values chosen were at least 10 grey values above background and autofluorescence in each case. Each binary mask was then processed: all single pixels were removed, and each binary mask was dilated by 5 pixels to ensure entire inclusion bodies were being identified. EM48+, EPR+, and MW1+ binary masks were combined to create an Htt inclusion body ‘master mask’. Each object within the master mask was considered a single Htt inclusion body.

To immunophenotype each Htt inclusion body, the maximum intensity of EM48, EPR, MW1, pan ubiquitin, K48-linked ubiquitin, K63-linked ubiquitin, ubiquilin 2, and p62 were measured within each object of the master mask. To determine whether an Htt inclusion body was positive or negative for each epitope, ubiquitin, or triage protein, intensity thresholds were manually determined for each case. Thresholds were set manually for each case to segment positive labelling only. Values chosen were at least 10 grey values above background and autofluorescence in each case.

To identify whether Htt inclusion bodies were nuclear or non-nuclear, we identified all nuclei; a binary mask of all Hoechst-positive nuclei was created using the Count Nuclei application. To identify whether each object in the master mask was nuclear, the maximum intensity of the nuclear binary mask within the master mask was also measured. An intensity measure of 1 denoted the inclusion was nuclear, whereas 0 denoted a non-nuclear inclusion.

### Statistical Analysis

Graphs were constructed in GraphPad Prism (version 9.3.1). Differences in the proportions of Htt inclusion body immunophenotypes between CAG repeat length, age of onset, and Vonsattel grade HD groups were tested using a chi-square (χ2) test for trend. Groups were determined based on the median for each HD clinicopathology measure; ≤43 or >43 for CAG repeat length, ≤40 or >40 for age of onset, and 1-2 or 3-4 for Vonsattel grade. As the data did not satisfy assumptions of normal distribution and equality of variance, a non-parametric Mann-Whitney test was performed to test for differences between normal and HD groups, and a non-parametric Wilcoxon test for paired samples was used to test for differences between the proportion of Htt inclusion bodies with positive or negative labelling for an epitope. A 2-way ANOVA with Šídák multiple comparisons test was used to test for differences in the mean Htt inclusion body size, percentage of nuclear inclusion bodies, and percentage of inclusion body phenotypes labelled for ubiquitin, K48- or K63-linked ubiquitin, or p62 or ubiquilin 2. Density plots for mean Htt inclusion body size were constructed in R (version 4.1.1) using the *ggplot2* package and the heatmap summarising the features of each Htt inclusion body epitope immunophenotype and ubiquitination and triage protein profile was constructed using the *ComplexHeatMap* package.

## Results

### Visualisation of Htt inclusion bodies in the HD brain through immunohistochemical profiling with Htt inclusion body epitope, ubiquitination, and triage protein antibodies

A multiplexed immunohistochemistry approach enabled the combined labelling of three previously validated Htt epitope antibodies (12) together with antibodies targeting specific ubiquitination patterns and associated triage proteins. The selection of the 3 Htt epitope antibodies were based on the demonstration of the highest number of aggregates captured with these antibodies in post-mortem HD cortical brain tissue (12). This approach allowed us to profile the pattern of Htt epitope co-labelling and the specific ubiquitination and triage protein profiles for each inclusion body.

The Htt antibodies EM48, EPR, and MW1 demonstrated varying degrees of co-labelling (Figure 1A-C). Inclusion bodies co-labelled with all three antibodies were often observed outside the Hoechst-positive nuclei (Figure 1A-C, I yellow arrow). The inclusion bodies observed within Hoechst-positive nuclei were frequently observed to co-label with EPR and MW1 antibodies (Figure 1A-C, I white arrow). The Htt inclusions also displayed immunoreactivity for antibodies targeting pan ubiquitination (Figure 1D) and specific K48- and K63-linked polyubiquitination (Figure 1E-F).

**Figure 1:**
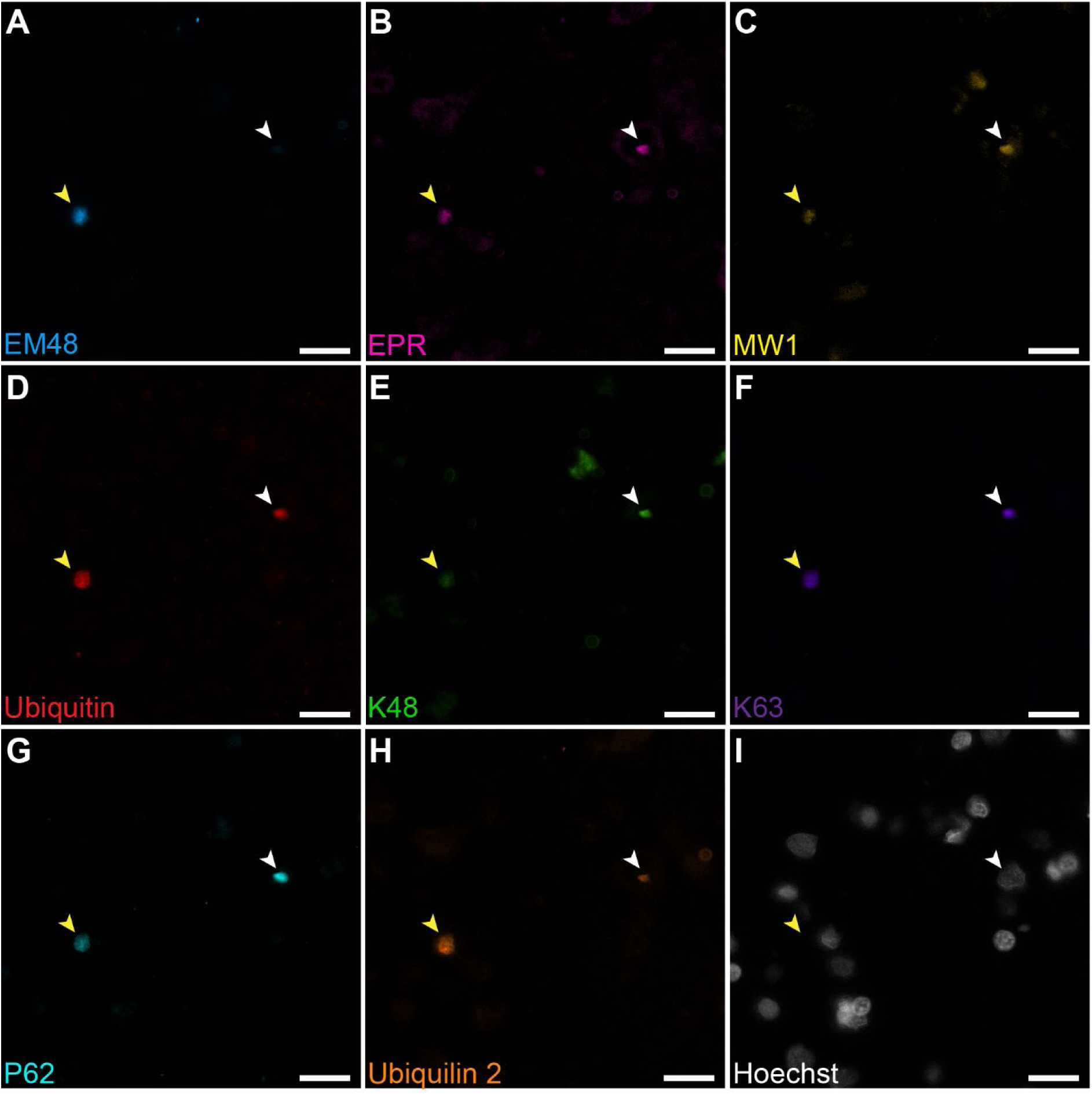
Immunohistochemical profiling of Htt inclusion body ubiquitination and associated triage protein binding in the HD human middle temporal gyrus. Multiplexed immunohistochemical approaches were used to identify Htt inclusion bodies, ubiquitin species, and triage proteins in neurologically normal and HD human middle temporal gyrus tissue microarray cores. Example images from HD case, HC150, are shown. Htt inclusion body antibodies, EM48 (A), EPR (B), and MW1 (C), were used for labelling together with antibodies for pan-ubiquitin (D), K48- and K63-linked polyubiquitination (E and F), p62 (G), and ubiquilin 2 (H), with a Hoechst nuclear counterstain (I); scale bars = 20 µm.

### Htt inclusion body antibodies against unique epitopes predominantly identify distinct Htt inclusion bodies in the human HD brain

In a previous study we validated Htt epitope antibodies using chromogenic single-label immunohistochemistry on separate tissue microarrays (12). Here, the use of multiplexed immunohistochemistry allowed for the characterisation of EM48, EPR, and MW1 within the same tissue microarray cores (Figure 2).

**Figure 2:**
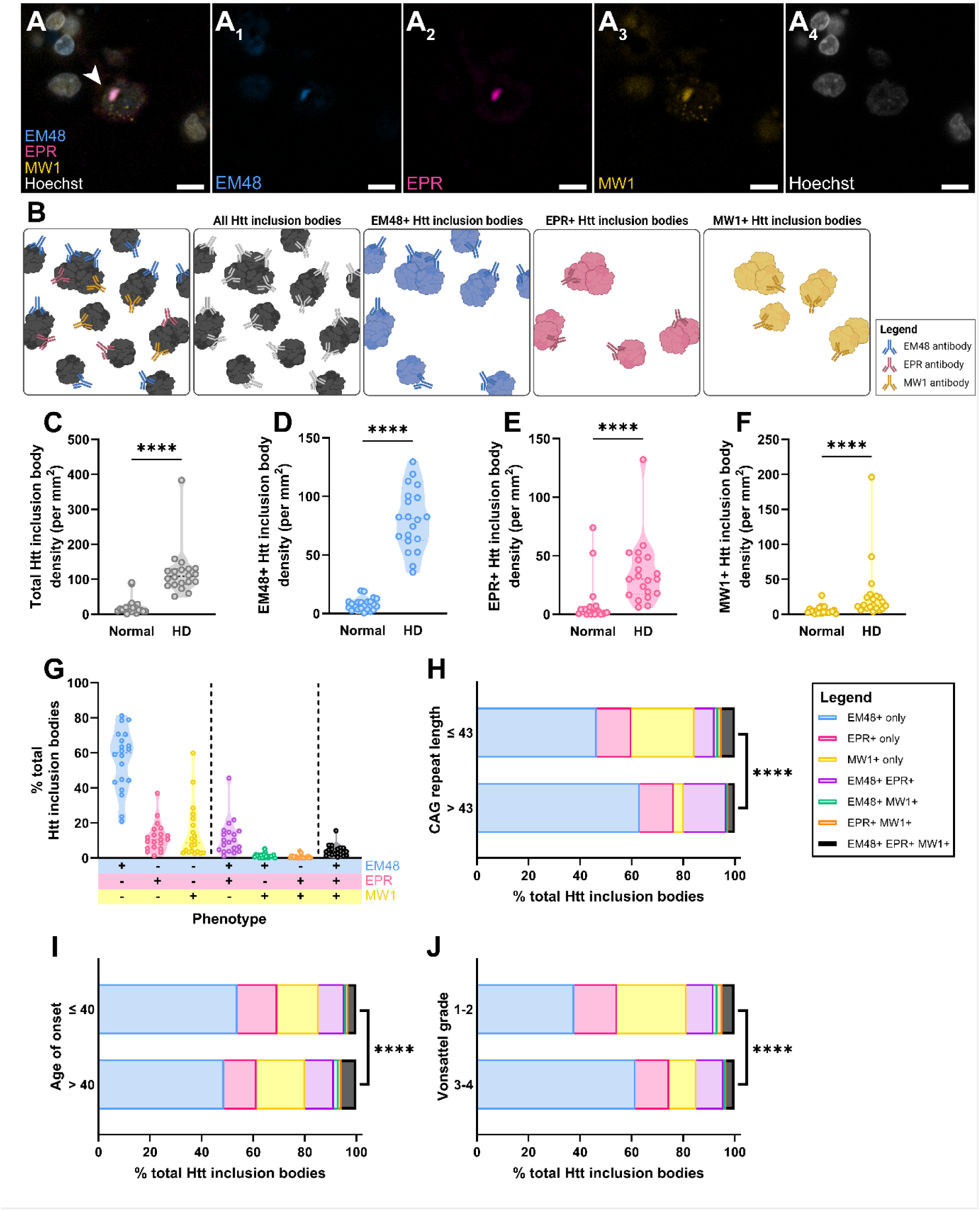
EM48+, EPR+, and MW1+ Htt inclusion bodies are increased in the HD middle temporal gyrus. Immunohistochemical labelling was used to visualise Htt inclusion bodies using EM48, EPR, and MW1 antibodies. A representative image of an EM48+ EPR+ MW1+ Htt inclusion body from HD case, HC145, is shown (A); scale bars = 10 µm. For quantification, total Htt inclusion bodies were identified by creating separate Htt inclusion body masks from EM48, EPR, and MW1 immunolabelling, which were combined to create an Htt inclusion body master mask. Each object within the Htt inclusion body master mask was considered a single Htt inclusion body and may be labelled with any combination of the three antibodies (EM48+, EPR+ or MW1+) (B). An inclusion body was classified with an EM48+, EPR+ or MW1+ phenotype if the maximum intensity for the respective immunolabel was above a manually determined threshold for each case (B). The density of total Htt inclusion bodies (C), EM48+ (D), EPR+ (E), and MW1+ (F) Htt inclusion bodies were compared between neurologically normal and HD cases using unpaired t-tests. Data presented as mean ± SD; normal n = 22 and HD n = 20. The percentage of Htt inclusion bodies with each possible combination of EM48, EPR, and MW1 labelling was calculated for each HD case (G). Data are presented as truncated violin plots (n = 20). The percentage of Htt inclusion bodies identified as each EM48, EPR, and MW1 +/- phenotype was compared between HD cases with a CAG repeat length of less than or greater than 43 (H), with an age of onset of less than or greater than 40 (I), and with a Vonsattel grade of 1-2 and 3-4 (J) using a Chi-square analysis. Data presented as stacked bar graphs representing all Htt inclusion bodies from all HD cases.

We first sought to validate that immunofluorescent labelling using EM48, EPR, and MW1 Htt epitope antibodies yielded the same Htt inclusion body density as previous chromogenic single-label approaches (12). We identified significant increases in EM48+, EPR+, MW1+ and total Htt inclusion body density in HD cases when compared with normal cases (Figure 2C-F). We subsequently determined the extent of co-labelling of Htt inclusion body epitope antibodies in HD cases by calculating the percentage of Htt inclusion bodies with each EM48, EPR, and MW1 +/- immunophenotype (Figure 2G). EM48+ only Htt inclusion bodies were the most abundant (56 ± 18%), followed by MW1+ exclusive (14 ± 15%), EPR+ exclusive (12 ± 8%), EM48+ EPR+ (11 ± 10%), EM48+ EPR+ MW1+ (4 ± 3%), EM48+ MW1+ (1 ± 1%), and EPR+ MW1+ (0.7 ± 1%). Therefore, on average, over 80% of Htt inclusion bodies were identified by only one epitope-specific antibody, with less than 20% containing two or more identifiable epitopes (Figure 2G).

To better understand the disease relevance of these Htt inclusion body immunophenotypes, we subsequently compared the percentage of Htt inclusion bodies with each EM48, EPR, and MW1 +/- phenotype in HD cases with a CAG repeat length of less than or greater than 43, with an age of onset of younger or older than 40, and with a Vonsattel grade of 1-2 (early pathology) and 3-4 (advanced pathology) (Figure 2H-J). We identified key differences in the proportion of Htt inclusion body immunophenotypes for all three disease characteristics. There was a higher relative proportion of inclusion bodies labelled with EM48+ only and corresponding lower relative proportion of MW1+ only and EM48+ EPR+ MW1+ inclusion bodies in cases with a CAG repeat length of greater than 43, an age of onset younger than 40, or advanced Vonsattel grades 3-4 (Supplementary Table 1). Therefore, the proportion of inclusion bodies with specific EM48, EPR, and MW1 +/- Htt immunophenotypes is associated with HD disease severity.

### EM48+ Htt inclusion bodies are smaller and predominantly non-nuclear in location, while EPR+ MW1+ inclusion bodies are larger and predominantly nuclear

We subsequently sought to further characterise the EM48, EPR, and MW1 +/- Htt inclusion bodies based on characteristics including: size and cellular location (Figure 3), ubiquitination patterns (Figure 4), and tagging with triage proteins (Figure 5).

**Figure 3:**
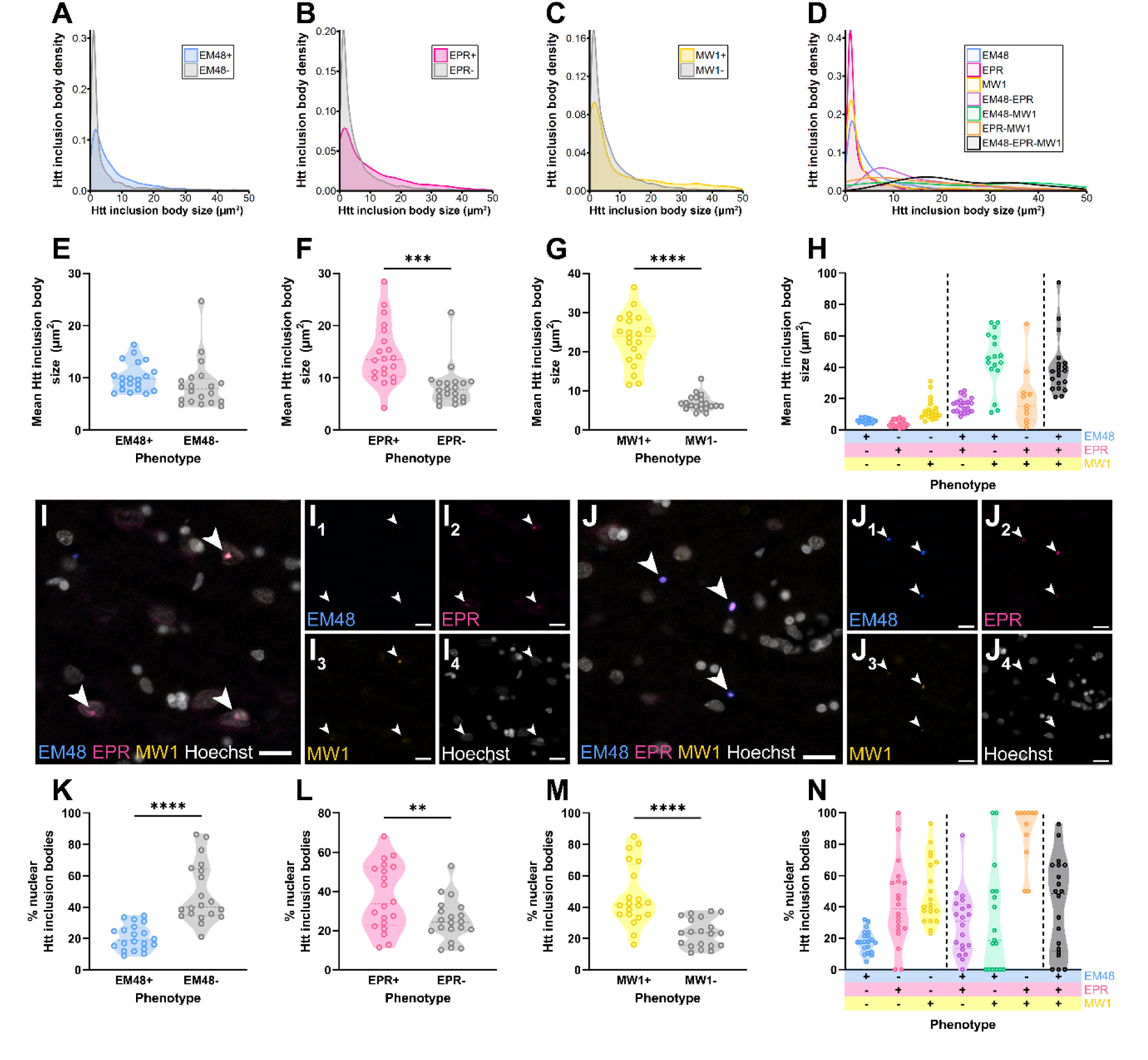
EM48+ only Htt inclusion bodies are smaller than other EM48-EPR-MW1 immunophenotypes and are predominantly non-nuclear in localisation. The size of each Htt inclusion body was determined and plotted as single-cell data on distribution graphs (A-D) and case-mean data on truncated violin plots (E-H, n = 20 cases) to compare the size between EM48+ and EM48-inclusions (A and E), EPR+ and EPR-inclusions (B and F), MW1+ and MW1-inclusions (C and G), and each EM48-EPR-MW1 phenotype (D and H). Htt inclusion bodies were noted to be both nuclear and non-nuclear; representative images of nuclear (I) and non-nuclear (J) Htt inclusion bodies from representative HD case, HC134, with EM48, EPR, and MW1 immunoreactivities are shown; scale bars = 20 µm. The percentage of EM48+ and EM48-inclusions (K), EPR+ and EPR-inclusions (L), MW1+ and MW1-inclusions (M), and each Htt inclusion body EM48, EPR, and MW1 phenotype identified within a nucleus was determined per HD case (n = 20). The mean Htt inclusion size and percentage of nuclear Htt inclusion bodies per case were compared between EM48+/-, EPR +/-, and MW1+/- inclusion groups using a Wilcoxon matched-pairs signed rank test and between each EM48, EPR, and MW1 +/- phenotype for HD cases using a mixed-effects analysis, with Geisser-Greenhouse correction and Tukey’s multiple comparisons test. Statistical significance of differences shown for E-G and K-M: ** p ≤ 0.01, *** p ≤ 0.001, **** p ≤ 0.0001. Statistical significance for H and N are shown in Supplementary Table 2.

**Figure 4:**
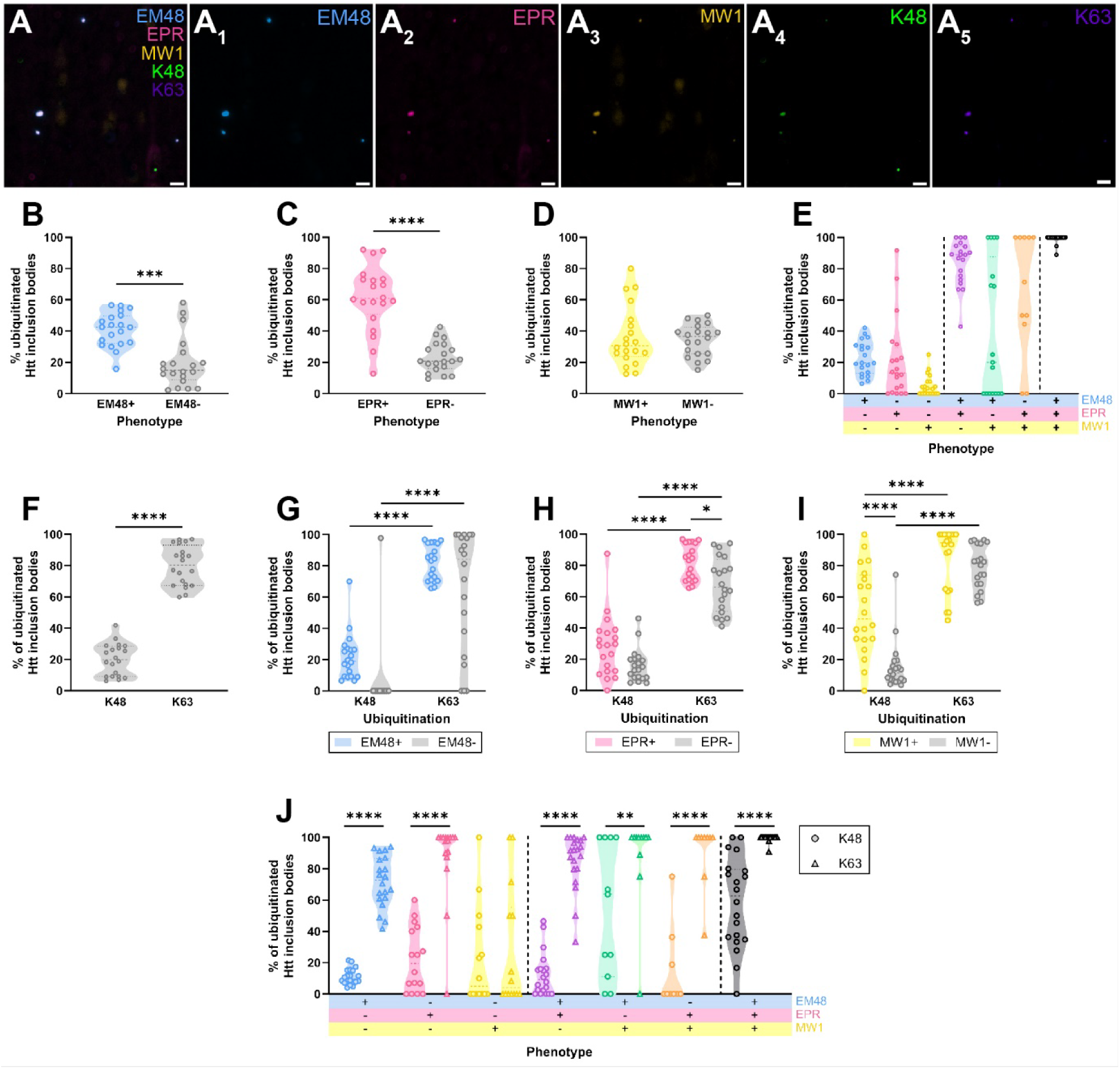
Htt inclusion bodies are not frequently ubiquitinated, but when ubiquitinated, are predominantly ubiquitinated by K63-linked ubiquitin. Immunohistochemical labelling revealed that EM48, EPR, and/or MW1 Htt inclusion bodies were ubiquitinated by K48- and/or K63-linked ubiquitin (**A**); a representative image of K48- and K63-ubiquitinated Htt inclusion bodies from HD case, HC145, is shown; scale bars = 10 µm. The ubiquitination status of each Htt inclusion body was determined by labelling for pan-, K48-, and K63-linked ubiquitin, where positive labelling was identified if the maximum intensity was above manually determined thresholds. The percentage of EM48+ versus EM48- (**B**), EPR + versus EPR- (**C**), and MW1+ versus MW1- (**D**) Htt inclusion bodies that were ubiquitinated (either pan, K48-, and/or K63-linked) were compared using a Wilcoxon matched-pairs signed rank test. The percentage of ubiquitinated Htt inclusion bodies was determined for each EM48, EPR, and MW1 +/- phenotype per HD case (**E**), and compared between phenotypes using a mixed-effects analysis, with Geisser-Greenhouse correction and Tukey’s multiple comparisons test. The percentage of ubiquitinated Htt inclusion bodies ubiquitinated by K48- versus K63-linked ubiquitin was compared using a Wilcoxon matched-pairs signed rank test (**F**). The percentage of ubiquitinated EM48+ versus EM48- (**G**), EPR+ versus EPR- (**H**), and MW1+ versus MW1-(**I**) Htt inclusion bodies ubiquitinated by K48-versus K63-linked ubiquitin were compared using an ordinary two-way ANOVA with Tukey’s multiple comparisons test. The percentage of EM48, EPR, and MW1 +/- immunophenotypes Htt inclusion bodies identified as being ubiquitinated by K48- or K63-linked chains were compared using an ordinary two-way ANOVA with Sidak’s multiple comparisons test (**J**). Data are presented as truncated violin plots (n = 20). Statistical significance of differences shown for B-D and F-J: * p ≤ 0.05, ** p ≤ 0.01, *** p ≤ 0.001, **** p ≤ 0.0001. Statistical significance for E is shown in Supplementary Table 3.

**Figure 5:**
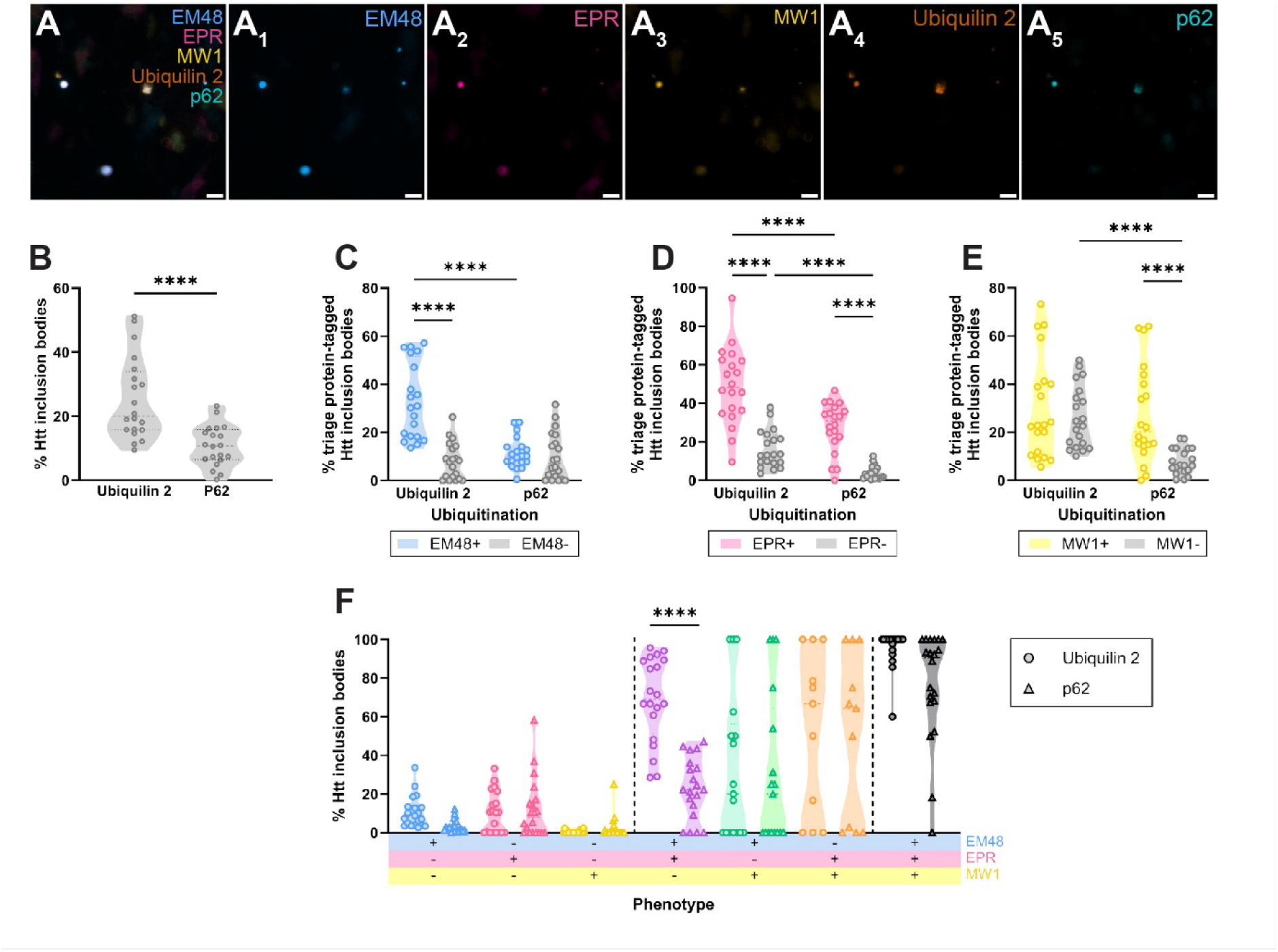
Ubiquitinated Htt inclusion bodies are more highly tagged by ubiquilin 2 than p62. Immunohistochemical labelling revealed that EM48, EPR, and/or MW1 Htt inclusion bodies were tagged by triage proteins, ubiquilin 2 and/or p62 (**A**); a representative image of ubiquilin 2- and/or p62-tagged Htt inclusion bodies from HD case, HC145, is shown; scale bars = 10 µm. The tagged triage protein status of each ubiquitinated Htt inclusion body was determined by labelling for ubiquilin 2 and p62, where positive labelling was identified if the maximum intensity was above manually determined thresholds. The percentage of ubiquitinated Htt inclusion bodies tagged by ubiquilin 2 versus p62 was compared using a Wilcoxon matched-pairs signed rank test (**B**). The percentage of EM48+ versus EM48- (**C**), EPR+ versus EPR- (**D**), and MW1+ versus MW1- (**E**) Htt inclusion bodies tagged by ubiquilin 2 or p62 were compared using an ordinary two-way ANOVA with Tukey’s multiple comparisons test. The percentage of EM48, EPR, and MW1 +/- immunophenotyped Htt inclusion bodies tagged by ubiquilin 2 or p62 were compared using an ordinary two-way ANOVA with Sidak’s multiple comparisons test (**F**). Data presented as truncated violin plots (n = 20 cases). Statistical significance of differences: *** p ≤ 0.001, **** p ≤ 0.0001.

We first sought to determine whether there were differences in the size of Htt inclusion bodies identified with EM48, EPR, and/or MW1 epitope-specific antibodies (Figure 3A-H). EM48+ inclusion bodies were not significantly different in size to EM48- inclusion bodies (Figure 3A and E, Supplementary Table 2), while EPR+ and MW1+ inclusion bodies were significantly larger than EPR- and MW1-inclusion bodies, respectively (Figure 3B, C, F, and G, Supplementary Table 2). When looking specifically at EM48, EPR, and MW1 +/- Htt inclusion body immunophenotypes, inclusion bodies identified by only one epitope-specific antibody (EM48+ only, EPR+ only, and MW1+ only) were generally smaller than those identified by two or more epitope-specific antibodies (EM48+ EPR+, EM48+ MW1+, EPR+ MW1+, and EM48+ EPR+ MW1+) (Figure 3D and H, Supplementary Table 2).

Previous research has described Htt inclusion bodies to be predominantly nuclear in location, however the presence of extranuclear inclusions has also been documented (13, 28, 47, 48) (Figure 3I). We observed EM48, EPR, and MW1 +/- Htt inclusion body immunophenotypes inclusion bodies that did not co-localise with Hoechst-positive nuclei (Figure 3J). We sought to determine whether there were differences in the nuclear versus non-nuclear distributions of Htt inclusion bodies identified with EM48, EPR, and/or MW1 epitope-specific antibodies (Figure 3K-N). A significantly lower percentage of EM48+ inclusion bodies were identified in the nucleus relative to EM48-inclusion bodies (Figure 3K, Supplementary Table 2). In comparison, a significantly higher percentage EPR+ and MW1+ inclusion bodies were identified in the nucleus relative to EPR- and MW1-inclusion bodies, respectively (Figure 3L and M, Supplementary Table 2). Upon examination of EM48, EPR, and MW1 +/- Htt inclusion body immunophenotypes specifically, a significantly lower proportion of EM48+ only inclusion bodies were nuclear in location compared with all other inclusion body phenotypes (Figure 3N, Supplementary Table 2). Therefore, Htt inclusion bodies labelled by EM48 are less likely to be located in the cell nucleus.

### Htt inclusion bodies immunoreactive for single epitopes are less ubiquitinated compared with Htt inclusion bodies immunoreactive for multiple epitopes

To understand if a particular Htt inclusion body immunophenotype was affiliated with certain protein degradation pathways, we sought to determine whether there were differences in the ubiquitination status of Htt inclusion bodies identified with EM48, EPR, and/or MW1 epitope-specific antibodies (Figure 4). We observed inclusion bodies immunoreactive for EM48, EPR and/or MW1 epitopes that also labelled for both K48- and K63-linked ubiquitin, or for K63- or K48-linked ubiquitin alone (Figure 4A). Overall, 59.9% of Htt inclusion bodies were ubiquitinated. A significantly greater percentage of EM48+ and EPR+ inclusion bodies were ubiquitinated (labelled for either pan-ubiquitin, K48- and/or K63-linked ubiquitin) relative to EM48- and EPR- inclusion bodies (Figure 4B-D, Supplementary Table 3). Upon examination of EM48, EPR, and MW1 +/- Htt inclusion body immunophenotypes specifically, a lower percentage of inclusion bodies identified by only one epitope-specific antibody (EM48+ only, EPR+ only, and MW1+ only) were ubiquitinated compared to inclusion bodies identified by two or more epitope-specific antibodies (EM48+ EPR+, EM48+ MW1+, EPR+ MW1+, and EM48+ EPR+ MW1+) (Figure 4E, Supplementary Table 3).

### Ubiquitinated Htt inclusion bodies are more likely to be poly-ubiquitinated by K63-linked than K48-linked ubiquitin

To further characterise the ubiquitination status of the Htt inclusion bodies, we investigated the frequency of specific ubiquitination by K63- and K48-linked ubiquitin chains (Figure 4). Of all Htt inclusion bodies that were ubiquitinated, a greater percentage (79.9 ± 12.6%) were immunoreactive for K63- compared to K48-linked ubiquitin (20 ± 10.2%, Figure 4F, Supplementary Table 3). Comparing Htt inclusion bodies classified as EM48, EPR, or MW1+/-, a significantly greater percentage of Htt inclusion bodies were ubiquitinated by K63- compared to K48-linked ubiquitin for all groups (Figure 4G-I, Supplementary Table 3). Similarly, the percentage of K63-linked ubiquitin-labelled Htt inclusions was significantly greater than K48-linked for all Htt inclusion body immunophenotypes, with the exception of the MW1+ only phenotype (Figure 4J, Supplementary Table 3). Therefore, Htt inclusions are more frequently ubiquitinated by K63- than K48-linked ubiquitin chains, irrespective of the Htt inclusion body immunophenotype.

### Ubiquitinated Htt inclusion bodies are more highly immunoreactive for ubiquilin 2 than p62

Lastly, we sought to determine whether ubiquitinated Htt inclusion bodies were differentially immunoreactive for triage proteins p62 or ubiquilin 2. We observed inclusion bodies immunoreactive for EM48, EPR and/or MW1 epitopes that also labelled for both ubiquilin 2 and p62, or for ubiquilin 2 alone (Figure 5A). A greater percentage of Htt inclusion bodies were immunoreactive for ubiquilin 2 than p62 (Figure 5B). A significantly greater percentage of EM48+ and EPR+ inclusion bodies were immunoreactive for ubiquilin 2 than EM48- and EPR- inclusion bodies (Figure 5C-D, Supplementary Table 4). In addition, a significantly greater percentage of EM48+ and EPR+ inclusion bodies were immunoreactive for ubiquilin 2 than for p62 (Figure 5C-D, Supplementary Table 4). However, MW1+ inclusion bodies demonstrated no difference in immunoreactivity for ubiquilin 2 but did show increased p62 immunoreactivity compared to MW1- inclusion bodies (Figure 5E, Supplementary Table 4). The inclusion bodies identified by two or more epitope-specific antibodies (EM48+ EPR+,

EM48+ MW1+, EPR+ MW1+, and EM48+ EPR+ MW1+) were more frequently immunoreactive for ubiquilin 2 and p62 compared to those identified by only one epitope-specific antibody (EM48+ only, EPR+ only, and MW1+ only) (Figure 5F, Supplementary Table 4). Notably, a significantly greater proportion of the EM48+EPR+MW1- immunophenotype inclusion bodies had ubiquilin 2 immunoreactivity than p62 (Figure 5F, Supplementary Table 4).

### Inclusion bodies recognised by multiple Htt antibodies have a distinct ubiquitination signature compared to inclusion bodies labelled with a single antibody

We finally sought to reconcile our data and determine whether different Htt inclusion body immunophenotypes have distinct signatures of ubiquitination and triage protein labelling (Figure 6). For each case we calculated the percentage of each phenotype that was nuclear in location, ubiquitin+, K48-linked ubiquitin+, K63-linked ubiquitin +, ubiquilin 2+ and p62+, along with the inclusion size. When plotted as a heatmap by Htt inclusion body immunophenotype, inclusion bodies had distinct sizes, and ubiquitination and triage protein signatures. EM48+MW1+EPR+ inclusion bodies were predominantly larger and more frequently ubiquitinated, K63-linked ubiquitin+, K48-linked ubiquitin+, ubiquilin 2+ and p62+. In comparison, the single-labelled inclusion bodies were smaller, less frequently nuclear, or ubiquitinated and not typically labelled for triage proteins. Also of note was the EM48+ EPR+ MW1- immunophenotype which had a similar signature to the triple-labelled inclusion bodies except for a lower frequency of K48-linked ubiquitin and p62 immunoreactivity. The EM48- EPR+ MW1+ phenotype was also interesting as it showed a predominantly nuclear location with variable degrees of ubiquitination and triage protein labelling between cases. These findings are visually summarised in Figure 6B.

**Figure 6:**
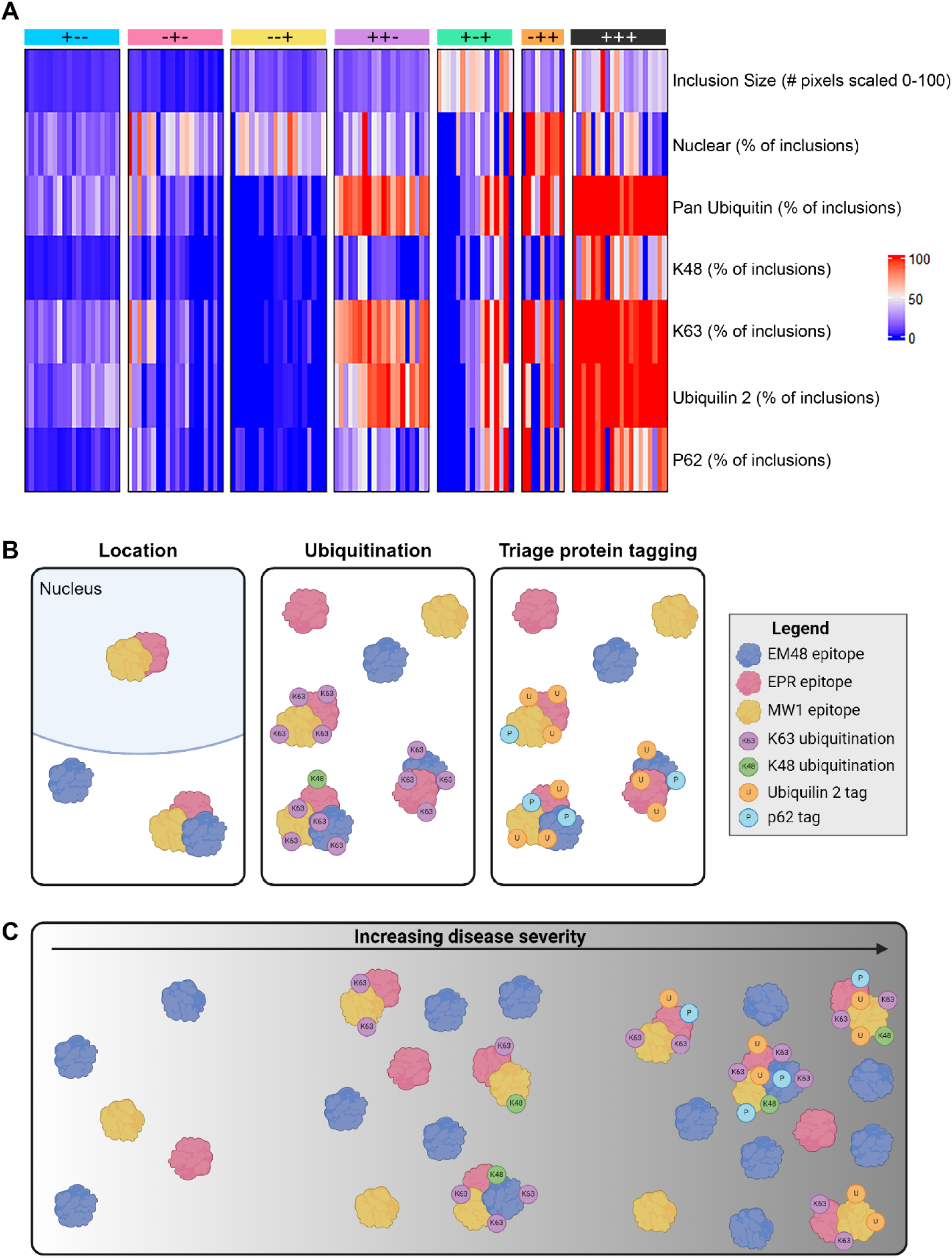
Summary of Htt inclusion body characteristics. Heatmap organised by Htt inclusion body phenotype, with each column representing a single case and each bar coloured according to that case’s value for the characteristic outlined by the row title **(A)**. Schematic illustrating the general characteristics of each Htt inclusion body phenotype: 1. EPR+MW1+ inclusion bodies are more frequently located in the nucleus compared to other phenotypes, 2. Htt inclusion bodies that label for more than one epitope-specific antibody are more frequently ubiquitinated, and that ubiquitination occurs more frequently by K63- compared to K48-linked ubiquitin chains, 3. Ubiquitinated Htt inclusion bodies are more frequently tagged by ubiquilin 2 than p62 **(B)**. Schematic summarising our hypothesis of Htt inclusion body immunophenotype, ubiquitination, and triage protein tagging with increasing HD severity **(C)**.

## Discussion

This study sought to investigate the ubiquitination profile of key Htt inclusion immunophenotypes immunolabelled with EM48, EPR5526, and MW1, which were previously validated to be HD-specific in human brain tissue microarrays (12). We immunohistochemically profiled the expression of ubiquilin 2 and p62 triage proteins known to bind to ubiquitinated proteins, as well as key ubiquitin linkages (K48 and K63) which target proteins for ubiquitination, in more than 50 cortical human brain samples from clinically and pathologically characterised HD and matching normal cases.

By co-labelling with EM48, EPR5526, and MW1 antibodies, which bind different epitopes of the Htt protein, we demonstrate the diversity in Htt inclusions present in the human brain and highlight the need to utilise a combination of antibodies to capture the full spectrum of inclusion pathology. This finding has implications for understanding the pathogenesis of HD, and for developing antibody-based therapeutic interventions. The heterogenous labelling of Htt inclusions implies that Htt inclusion composition and/or conformational properties is diverse and raises the question of how inclusions with different immunophenotypes contribute to the neurodegenerative processes. Based on the immunophenotyping, and relationship to HD clinicopathology, and ubiquitination and triage protein tagging signatures, we have formed a hypothesis of how these different Htt inclusion bodies form with increasing disease severity shown in Figure 6C.

Inclusions are an accumulation of different aggregated forms of Htt proteins or Htt aggregates bearing different patterns of PTMs, and the heterogeneity of immunophenotypes likely reflects this composition. EM48 binds a sequence that spans the 256 amino acids encompassing the N-terminus, polyQ and PRD regions, while MW1 and EPR bind the most distal N-terminal sites (49–51). The differential binding of these antibodies likely reflects the diversity in (1) the conformational properties of Htt in different aggregated forms, (2) the presence of distinct PTMs and patterns, and/or (3) the biochemical composition (e.g. proteins and lipids) and interactome of the Htt aggregate within inclusions. All of these factors could contribute to the differential labelling of Htt inclusions by antibodies targeting different epitopes. There are various PTMs that occur of amino acids within the N-terminal EPR and MW1 epitopes, but whether these antibodies bind irrespective of such modifications is unknown (52). Studies of recombinant Htt protein are needed to elucidate the specificity of antibody binding in the presence of post-translational modifications and truncations.

The results from our investigation also reinforce that the size, structure, and spatial localisation of Htt inclusions likely influences the amount of labelling by antibodies targeted to different epitopes. A study of Htt inclusion formation using an in-vitro model revealed that cytoplasmic and nuclear inclusions have distinct ultrastructural properties (53). Mature cytoplasmic Htt inclusions have a core and shell architecture that sequesters a variety of lipids, proteins and organelles. Whereas nuclear inclusions are more fibrillar and do not have a core and shell organisation. These structures remain to be captured and validated using in-vivo models or post-mortem human tissue studies but provide key insights into Htt inclusion heterogeneity. Electron microscopy of the Httex1 72Q inclusions in HEK 293 cells showed that while the outer shell of cytoplasmic inclusions is densely labelled by Htt antibodies, the inner core is tightly packed with minimal labelling (53). Therefore, the size, structure, and location of Htt inclusions likely influences the degree of labelling by antibodies targeted to different epitopes.

Our results also demonstrate that double- and triple-labelled inclusions were larger in size, with inclusions immunolabelled for either MW1 or EPR epitopes being significantly larger than those without. This finding indicates that EM48 preferentially binds relatively small and possibly less compact inclusions. Our finding of EM48 only inclusions being predominantly non-nuclear corroborates previous studies which demonstrated EM48+ inclusions are mostly localised to the neuropil and cytoplasm of neurons, with some, but few present in the nucleus (13, 54, 55). The increased proportion of EM48 only inclusions in HD cases with greater CAG repeat length, higher Vonsattel grade, and earlier age of onset suggests that these predominantly small, non-nuclear inclusions are more associated with increased disease severity.

Notably, EM48-EPR+MW1+ inclusions were the only immunophenotype that was predominantly nuclear. This finding suggests that the distinct structure of nuclear inclusions may affect EM48 antibody binding. Of the multi-labelled phenotypes, the EM48+EPR+MW1- inclusions were the most abundant. These inclusions were the smallest of the multi-labelled inclusions and were predominantly non-nuclear. Previous studies have also documented the occurrence of cytoplasmic inclusions and other non-nuclear inclusions in HD post-mortem tissue (56, 57). Our data also indicated that the number of small non-nuclear EM48 only inclusions were associated with increased disease severity, however, these inclusions were less frequently ubiquitinated with either pan, K48-, and/or K63-linked ubiquitin. This finding potentially implies the cell may not be actively trying to remove these EM48 only inclusions by autophagy or the UPS, which are both reliant on ubiquitination as a signal for these protein degradation pathways. Therefore, we sought to investigate the ubiquitination patterns for the various inclusion phenotypes in more detail.

Our data demonstrates that inclusion bodies which label for multiple epitopes are more frequently ubiquitinated. These ubiquitinated inclusion bodies are more frequently ubiquitinated by K63- rather than K48-linked ubiquitin, irrespective of the Htt inclusion body phenotype, suggesting that autophagy is the primary mechanism by which the cell tries to remove large inclusions in HD. Previously, rodent studies have demonstrated that ubiquitination by K48-linked ubiquitin promotes Htt degradation, while ubiquitination by K63- linked ubiquitin promotes Htt aggregation (27). An age-related reduction in the K48-specific E3 ligase, Ube3a, is thought to drive an increase in K63-linked ubiquitination and subsequent formation of inclusions. While we did not investigate Ube3a levels in this study, our finding of predominant K63-linked ubiquitination in post-mortem human HD tissue supports this hypothesis. Overexpression of Ube3a in HD rodent models results in increased K48-linked ubiquitin-mediated degradation through the UPS, along with reduced K63-linked ubiquitination and Htt aggregation. Therefore, switching Htt ubiquitination from K63- to K48- linkage could be a promising therapeutic strategy to reduce mutant Htt without concurrently reducing normal Htt.

We investigated co-labelling of mHtt with the triage proteins ubiquilin 2 and p62 which target ubiquitinated proteins to the UPS or autophagy pathways. Previous studies demonstrated that mHtt inclusions impair the protein degradation process via inhibition of the UPS (58). Furthermore, observations of Htt-enriched structures resembling autophago-lysosomal vesicles in the HD brain led to the hypothesis that autophagy becomes dysfunctional in HD, resulting in insufficient Htt aggregate removal (57, 59). Our data showed that mHtt inclusions identified by two or more epitope-specific antibodies were predominantly co-labelled with ubiquilin 2, rather than p62. This observation is in line with observations by others that ubiquilin proteins (1, 2, 4) can be sequestered by Htt inclusions (60). Recent studies undermine the hypothesis of UPS disruption by Htt inclusions. Evidence of the dynamic relationship between the UPS and Htt aggregates, with UPS components transiently interacting with aggregates, demonstrates that the UPS is not prevented from carrying out its function as a result of being sequestered within aggregates (61). Future functional studies would need to explore whether mechanisms involved in protein sequestration and autophagy are impaired in cells with these inclusions.

### Limitations of study

There are several methodological considerations for our study. The use of tissue microarrays provides a high-throughput screening platform to examine Htt inclusion heterogeneity and ubiquitination. The advantage of this approach is experimental standardisation and efficiency, whereby all cases are labelled and imaged with the same conditions (39). However, the relatively small 2-mm diameter cores give a limited tissue area from each case to study. While the samples included in the tissue microarray include cortical layers II-V, larger tissue sections encompassing all cortical layers would be required to examine any layer-specific differences in inclusion immunophenotypes.

Our study also focused on the MTG which is one of the least degenerated regions of the cortex in HD (6). This region gives us insight into the earlier pathogenic processes involved in Htt accumulation based on the known presence of aggregates and mild neuronal loss, but it is not a region typically associated with HD symptomatology. Future studies would need to explore the relationship between Htt inclusion immunophenotype and ubiquitination profile in cortical regions more correlated with HD symptomatology.

Lastly, we used multiplexed labelling to examine the co-expression of several proteins of interest on the same tissue section, which was essential to answer our biological questions. However, a limitation of this method is that it relies on widefield fluorescence imaging to capture the images of five fluorophores simultaneously and accurately register images between labelling rounds. Future studies that use higher resolution imaging techniques would be useful to examine the relationship between ubiquitination and Htt inclusion ultrastructure in more detail.

## Conclusion

In conclusion, our study has demonstrated that Htt inclusion bodies have a diverse array of immunophenotypes based on co-labelling by three antibodies that bind different epitopes of the protein. We found that Htt inclusions with different immunophenotypes have distinct molecular profiles in terms of size, subcellular location, and ubiquitination. Most notably we found that small inclusion bodies labelled only by EM48 were predominantly non-nuclear, and non-ubiquitinated and their frequency was associated with disease severity, suggesting possible involvement in disease progression. Conversely, we found that large, multi-epitope labelled inclusion bodies are predominantly ubiquitinated by K63-linked ubiquitin, indicating a preferential targeting for degradation by autophagy. Our findings highlight how a combination of antibodies is required to capture the full spectrum of Htt pathology in Huntington’s disease and future studies should consider the diversity of inclusion body composition in the development of antibody-based therapeutic interventions.

## Abbreviations

CAG: glutamine
HD: Huntington’s disease
Htt: Huntingtin protein
LC3: microtubule- associated protein 1 light chain 3
MTG: middle temporal gyrus
PB1: Phox and Bem1 domain
PMD: post-mortem delay
PTM: post-translational modification
TMA: tissue microarray
UPS: ubiquitin-proteosome system.

## Declarations

### Ethics approval and consent to participate

The tissue was donated with written informed consent from the donor and family before brain removal. All research procedures and protocols were approved by the Health and Disability Ethics Committee (Ref.14/NTA/208).

### Consent for publication

Written informed consent to publish anonymised data pertaining to tissue donors was obtained from next of kin at the time of tissue donation. All research procedures and protocols were approved by the Health and Disability Ethics Committee (Ref.14/NTA/208).

### Authorship contribution statement

**Molly E.V. Swanson:** Writing – review & editing, Writing – original draft, Visualization, Validation, Software, Methodology, Investigation, Formal analysis. **Adelie Y.S. Tan:** Writing – review & editing, Methodology. **Lynette J. Tippett:** Writing – review & editing, Resources, Methodology. **Clinton P. Turner:** Writing – review & editing, Resources, Methodology. **Maurice A. Curtis:** Writing – review & editing, Resources, Methodology, Funding acquisition. **Emma L. Scotter:** Writing – review & editing, Resources, Methodology. **Hilal A. Lashuel:** Resources, Methodology. **Mike Dragunow:** Writing – review & editing, Resources, Methodology, Funding acquisition. **Richard L.M. Faull:** Writing – review & editing, Resources, Methodology, Funding acquisition. **Helen C. Murray:** Writing – review & editing, Writing – original draft, Visualization, Validation, Software, Methodology, Investigation. **Malvindar K. Singh-Bains:** Writing – review & editing, Writing – original draft, Resources, Project administration, Methodology, Investigation, Funding acquisition, Conceptualization.

## Acknowledgements

We express our appreciation to all donor families in New Zealand, who, through their generosity, have enabled their invaluable tissue donation to the Neurological Foundation Human Brain Bank for the construction of human brain TMAs. We thank Senior Histologists S. Amirapu and F. Biggins for contributing to tissue processing and paraffin embedding protocols for TMA preparation. We acknowledge the excellent work and assistance of M. Eszes (Human Brain Bank), K. Hubbard, R. Parker, C. Webb-Robinson, and E. McShane (Research Technicians). We acknowledge the staff at the Biomedical Imaging Research Unit for VSlide Scanner support.

## Data availability

The authors confirm that all other data supporting the findings in this study are available within the article and its Supplementary Material. Raw data will be shared by the corresponding author upon request.

